# The amino acid composition of a protein influences its expression

**DOI:** 10.1101/2023.03.28.534610

**Authors:** Reece Thompson, Benjamin Simon Pickard

## Abstract

The quantity of each protein in a cell only is only partially correlated with its gene transcription rate. Independent influences on protein synthesis levels include mRNA sequence motifs, amino acyl-tRNA synthesis levels, elongation factor action, and protein susceptibility to degradation. Here we report two novel forms of interaction between the amino acid composition of a protein and its expression level.

In animals, the differing origins of amino acids define a nutritional classification system and indicate their potential for scarcity – essential amino acids (EAA) are solely obtained from dietary supply, non-essential amino acids (NEAA) from biosynthetic supply, and conditionally essential amino acids (CEAA) from both. Accessing public proteomic datasets, we demonstrate that CEAA sequence composition is inversely correlated with expression – a rule of supply that is further magnified by rapid cellular proliferation. Similarly, proteins with the most extreme compositions of EAA are reduced in abundance. Homeostatic responses to malnutrition may result from the reductions in expression of extreme composition proteins participating in biological systems such as taste and food-seeking behaviour, oxidative phosphorylation, and chemokine function. The rule can also influence general human phenotypes and disease susceptibility: stature proteins are enriched in CEAAs, and a curated dataset of over 700 cancer proteins is significantly under-represented in EAAs.

A second rule, whereby individual amino acids influence protein expression is also described. This rule is shared across all kingdoms of life and rooted in the immutable structural and encoding parameters of each amino acid. Species-specific environmental survival pathways are shown to be enriched in proteins with amino acid compositions favouring higher expression according to this rule.

These two rules of protein expression regulation promise new insights into systems biology, evolutionary studies, experimental research design, and public health intervention.

## Introduction

The regulated transcription of mRNA from DNA, and subsequent translation into effector proteins, underlies all of life’s dynamic processes. However, a typical gene’s levels of mRNA and protein only show a correlation of 0.6 [1–3], indicating the presence of DNA-independent regulatory influences on translation. Those influences are complex and incompletely understood [4–6] but include mRNA sequence motifs, compatibility between mRNA codon choice and corresponding tRNA-amino acid availability [7, 8] and the complex regulation of translation initiation, elongation and termination.

The established mTORC1 signalling pathway elicits molecular and cellular changes in response to nutritional state via the monitoring of certain amino acid concentrations [9]. However, the direct impact of global amino acid scarcity on protein translation is underexplored, despite supply characteristics defining an important amino acid classification system in animals [10]. That classification comprises *essential* amino acids (EAA) required from diet, *non-essential* amino acids (NEAA) obtained through biosynthesis, and an ill-defined intermediate class, *conditionally essential* amino acids (CEAA), requiring supplementation from diet during development and periods of stress or illness [11, 12]. Over 500 million years ago the new animal kingdom was, in part, distinguished by a coordinated inactivation of biosynthetic pathways for the EAA class [13–15]. The resulting switch from an autotrophic to auxotrophic lifeway obliged the direct or secondary sourcing of EAAs from a diet of prototrophic plants, or heterotrophic prey that fed on those plants. The opportunity for increased biological complexity offered by the energetic efficiency of a higher trophic level has been of demonstrable advantage to animals but it created vulnerability to situational deficits in dietary supply of EAA and possibly CEAA. Such deficits would likely act by limiting tRNA-amino acid synthesis and availability, slowing the protein translation rate, and decreasing protein expression – with inevitable phenotypic consequences.

Here we present evidence obtained from quantitative proteomics datasets that a protein’s amino acid composition correlates with its expression in two ways. Firstly, we show that the proportions of the three nutritional classes of amino acids in an animal protein exert an influence on expression reflecting extrinsic supply and intrinsic amino acid biosynthesis constraints, respectively. A second form of intrinsic amino acid composition effect on expression is also described that is shared across all kingdoms of life and derived from the universal structural and encoding parameters of amino acids. We propose that evolution has harnessed both the extrinsic and intrinsic effects to select protein compositions that confer advantageous expression responses during environmental stress. These two rules offer intriguing new insights into the environmental influences on protein evolution and expression regulation.

## Results

### Extrinsic effects of amino acid classes on protein expression

We first examined how the amino acid nutritional class composition of every human protein influences its expression. Mass spectrometry-derived expression levels of 9,399 liver proteins were accessed from the public proteomics repository, PaxDB [16] (**Methods**). **Fig 1a** shows proteins ranked from low-to-high frequency of each of the three nutritional classes plotted against a conservative moving median protein expression level. A greater compositional frequency of CEAA (fCEAA) was generally associated with a modest decrease in protein expression, whereas greater EAA (fEAA) was associated with increased expression. However, at the extremes of composition, the outcome was more complex with high fEAA and fCEAA both repressing expression, and low values releasing constraints on expression. At both extremes, we interpret the apparent fNEAA influence on expression to be merely the passive consequences of active fCEAA or fEAA constraint effects. These findings are striking in two regards: they are naïve of mRNA expression information, and data are derived from human donors without known dietary amino acid deficiency.

**Figure 1.**
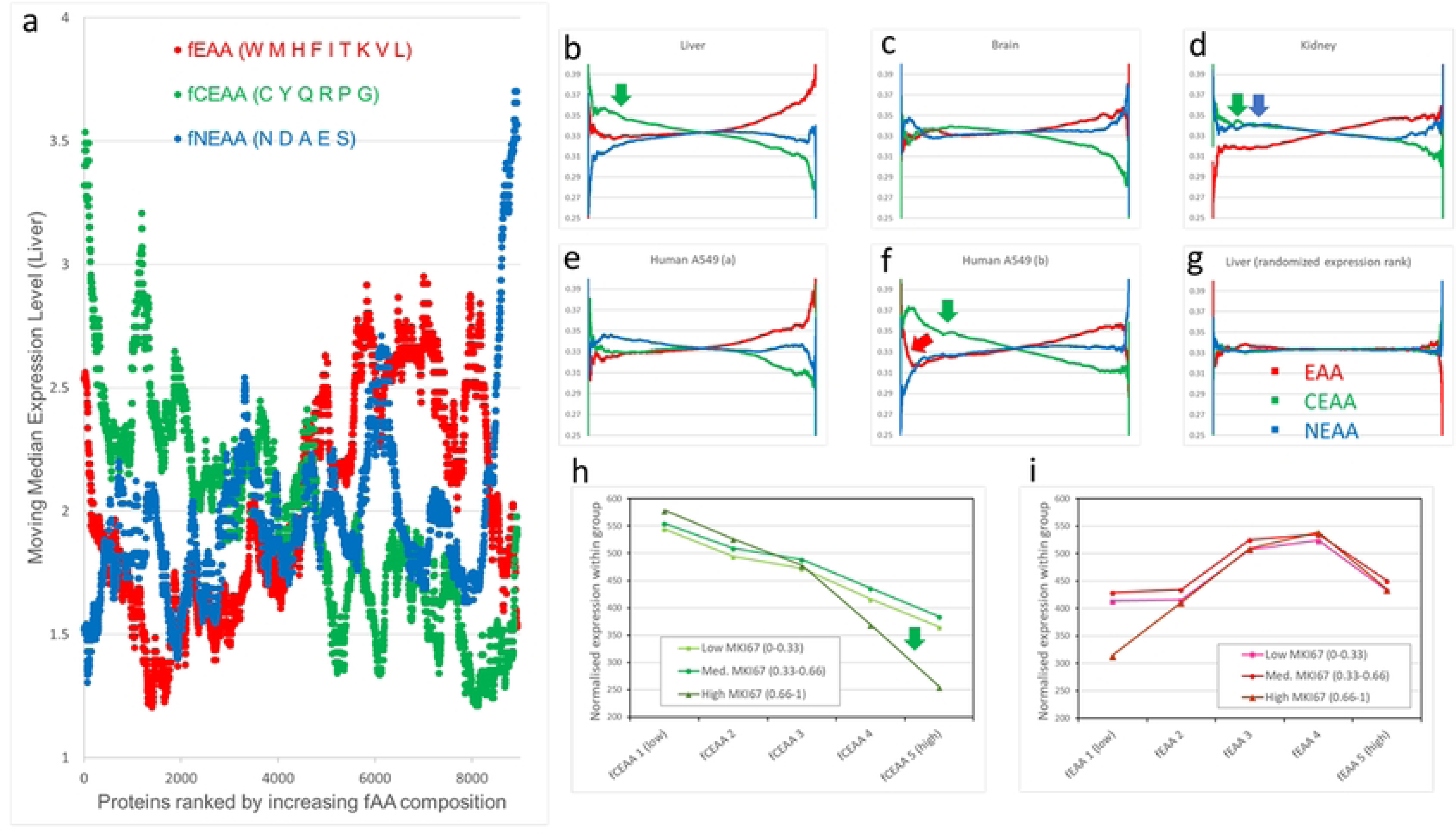
The relative proportions of the three nutritional amino acid classes change with protein expression level and proliferation rate. (a) Proteins were ranked by frequency of each of the three nutritional classes (NEAA, blue; CEAA, green; EAA, red. Single-letter amino acid codes are shown) and plotted against a moving median of liver expression levels (periodicity = 5% of total protein number) to determine influence on expression. (b-g) A smoothing procedure (see Methods and Supplementary File 1) was applied to visualise trends in relative, ranked amino acid class proportion when plotted against ranked protein expression level in human liver, brain, and kidney, and in A549 lung cancer cells lines with different proliferation rates (data from PaxDB). The full set of human organ data is presented in Supplementary Figure 1. In (g), liver proteins have been randomised with respect to expression rank, removing trends in amino acid class representation and confirming the validity of the smoothing approach. (h/i) The impact of cellular proliferation rate on protein expression was examined in PaxDB data from 26 cell lines from multiple laboratories stratified into three cohorts (low proliferation rate, 11 lines; medium, 8 lines; high, 7 lines) via normalised expression level of the proliferation marker MKI67 (Ki67). Proteins were additionally subdivided into 5 amino acid class frequency ranges (fEAA/fCEAA 1-5, with 5 having the greatest representation of the amino acid class). Normalised total protein expression levels were calculated for each of the 15 groups and plotted.

An alternative approach to data visualisation, focusing on relative amino acid composition, was applied to data from eight human organs and a lung alveolar basal epithelial adenocarcinoma cell line, A549. The accessed organ expression datasets (PaxDB) had already been normalised and integrated from several analyses. Ranked protein expression levels were plotted against smoothed relative EAA:CEAA:NEAA proportions for each protein (**Methods, Fig 1b-g**, and **Supplementary Fig 1**). The right side of every component image is largely conserved in appearance: high-level protein expression requires amino acid composition to be near to the population means (fEAA, 0.41; fCEAA, 0.28; fNEAA, 0.31). By contrast, the left side of each image was observed to fall into one of three distinct profiles. In the first, **Fig.1b, Supplementary Fig 1** liver, heart, and male and female gonads show a marked increase in proteins with high CEAA proportion at lower expression levels (green arrow). This suggests the existence of a biosynthetic shortfall of these amino acids that results in reduced expression of such proteins. In **Fig 1c, Supplementary Fig 1 p**ancreas and brain tissues do not appear to be affected by the steady-state supply of amino acids. In **Fig 1d, Supplementary Fig 1** kidney and lung CEAA and NEAA biosynthesis are seemingly both constrained (green and blue arrows) causing a greater proportion of those amino acids in proteins with low expression, with lower EAA levels reflecting this passively. These three profiles may reflect inherent organ features such as non-proteogenic amino acid use (gluconeogenesis), local proliferation rate, basal metabolic rate, or amino acid transportation capacities.

In **Fig 1e and f**, markedly different profiles are shown for two expression datasets from the same lung cancer cell line, A549. We hypothesised that differing tissue culture protocols in the source laboratories affected amino acid availability. In **Fig 1e** (A549 a, data from [17], #id: 3312331274) there is little evidence for constrained protein expression. However, in **Fig 1f** (A549 b, data from [18], #id: 878737823) - with cells described as actively proliferating - a substantial increase in CEAA and EAA proportions (green and red arrows) was observed at lower expression levels indicating that proteins with greater proportions of those two amino acid classes were inefficiently translated. A suspected interaction between proliferation rate, nutritional class and expression was therefore examined in PaxDB expression data from 26 cell lines. Wide provenance, intrinsic cell line expression differences, and uncertain culture conditions at the time of protein isolation required an objective means to stratify cell line data by proliferation rate. A normalised protein expression level was derived for established proliferation marker Ki67 (MKI67) within each cell line (this value correlated well with expression of cell division proteins such as CKS2, KIF23, POLA1, CDC45 and SPC24; data not shown). Cell lines were stratified into 3 groups by this proliferation rate proxy, as well as into 5 fEAA or fCEAA classes. When total protein expression levels within each of the resulting fifteen subdivisions were summed and plotted (**Fig 1h and 1i**), we saw evidence for expression influenced by two forms of amino acid supply kinetics. Firstly, a proportionately negative influence on expression exists across the full range of fCEAA, which is further intensified (green arrow) by the amino acid demands of rapid proliferation (**Fig 1h**). Secondly, a largely proliferation-independent, positive effect of increasing fEAA on protein expression levels was observed which switches to negative only for those proteins with the very highest fEAA (perhaps determined by the limits of EAA availability in media and its cellular uptake) (**Fig 1i**).

### Intrinsic effects on protein expression

We looked beyond nutritional supply constraint effects and amino acid groupings to determine if fundamental effects on protein expression existed at the individual amino acid level. Multiple Linear Regression (MLR) was used to determine individual amino acid frequency effects on expression within eight human organs and, in parallel, the root of plant *Arabidopsis thaliana* (a prototroph), the fungus *Saccharomyces cerevisiae*, and bacterium *Escherichia coli* (both heterotrophic/autotrophic) (**Table 1, Supplementary File 1**). The individual amino acid effects on expression were substantially conserved in scale and direction between human organs and across species. Increased representation of amino acids lysine (K), glycine (G), alanine (A), and valine (V) largely correlated with increased expression of a protein, whereas tryptophan (W), arginine (R), cysteine (C), and serine (S), largely correlated with reduced expression. Human-specific inhibitory effects were seen for isoleucine (I) and proline (P), and methionine (M) was consistently inhibitory in non-animal species. Aspartate (D) only showed a positive influence in humans. These MLR models are statistically highly significant, but only generate modest r^2^ values within a range of 0.02 to 0.13 for global prediction of protein expression in the absence of mRNA expression data or nutritional deficiency. We noted that the largest r^2^ values were observed for human ovary, heart, and testis, all represented within the subgroup represented in **Fig. 1b** and **Supplementary Fig. 1a**, and for the two microorganisms cultured in proliferation-driving conditions. This suggested that there may be overlapping effects from intrinsic amino acid characteristics and extrinsic amino acid constraint influences. This found further support in the over-representation of CEAAs within the group of amino acids showing MLR negative effects. Surprisingly, Serine (a NEAA not expected to be limiting) was also negatively associated with expression level. Serine’s vital role as a carbon source for the synthesis of other amino acids, and the pathological consequences of its deficiency in humans has prompted a recent call for its reclassification as a CEAA [19].

**Table 1.**
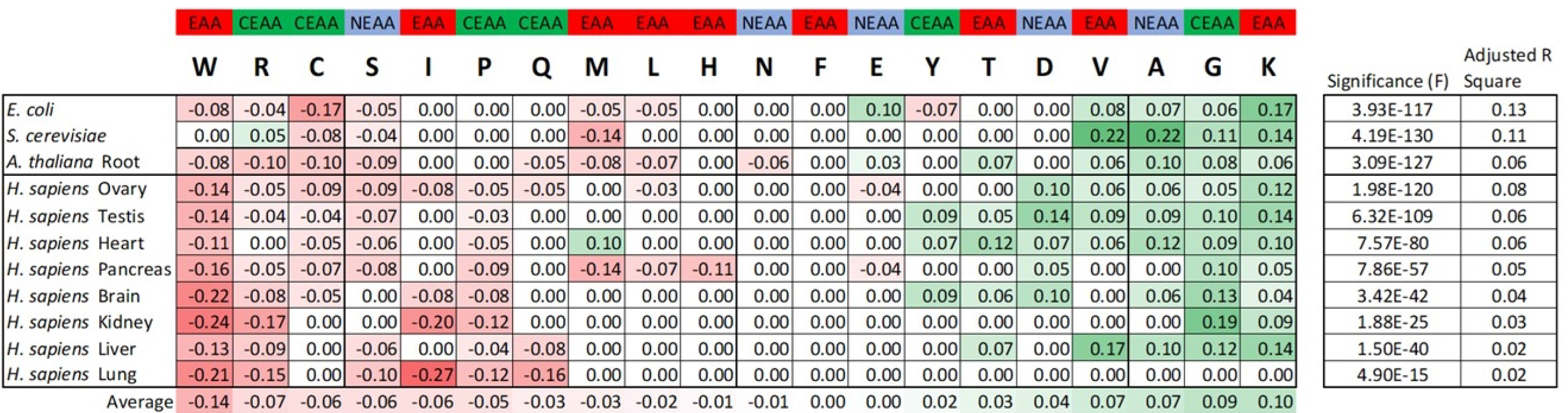
Pan-species conservation of individual amino acid influences on protein expression levels. Human organ data are shown in the lower section of the table and species data in the upper section. Numbers in cells represent the normalised magnitudes for statistically significant multiple linear regression (MLR) coefficients for each amino acid in each species or organ. Amino acids have been ordered left-to-right across the table from greatest average negative effect (shades of red) to greatest average positive effect (shades of green) on protein expression. Human amino acid nutritional class assignments are shown at the top of the table. Adjusted correlation values and statistical significance are shown on the extreme right of the table for each of the 11 MLRs. See **Supplementary File 1** for full MLR data.

The pan-species nature of these amino acid effects was effectively demonstrated by using the *E. coli* MLR model from **Table 1** and **Supplementary File 1** to predict trends in global human protein expression (**Fig. 2**). A moving average expression describing three orders of magnitude was observed when the liver MLR model value for each protein was plotted against that protein’s true liver expression (**Fig 2a**). The bacterial MLR model applied to human liver expression generated was still able to describe a moving average expression spanning two orders of magnitude (**Fig 2b**). To explain these universal effects of amino acids on protein expression we considered three fundamental properties as candidate influences. The first property was the number of synonymous codons assigned to each amino acid, in what we hypothesised was a proxy for the effect of codon choice on translation. Secondly, three related models of amino acid biosynthetic cost were applied to determine if metabolic economisation has directed protein evolution towards ‘thrifty’ amino acid composition and protein expression levels. In the Akashi [20] and Wagner models [21], the cost of synthesis for each amino acid is measured as high-energy phosphate bond equivalents and, in the Zhang model [22], this is further refined by including amino acid degradation constants. The third assessed property was the composite ‘Dufton score’ [23], assigned to amino acids based on their spatial volume and chemical complexity - encapsulating biosynthetic, structural, and functional parameters.

**Figure 2.**
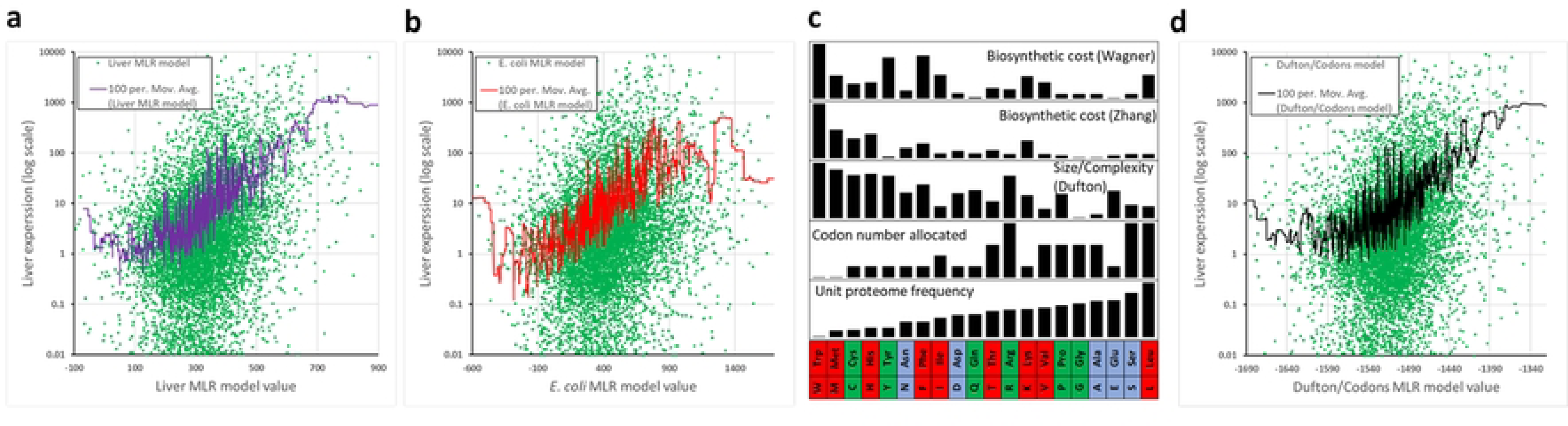
Pan-species effects of amino acids on protein expression can be largely explained by two fundamental parameters. The two expression prediction models generated by MLR analysis of individual amino acid effects on human liver and E. coli protein expression levels (detailed in **Table 1** and **Supplementary File 1**) were tested for their ability to predict global liver protein expression. Each individual protein is shown as a green dot representing model predicted expression and actual human liver expression (log scale). Model ability is visualised by plots of moving averages (purple, red, black: periodicity of 100 proteins). (**a**) Liver model on liver expression analysis. (**b**) E. coli model on liver expression analysis. (**c**) Individual properties of amino acids (identified by their the one- and three-letter designations at the bottom) are visualised along with their amino acid frequency in the unit human proteome, and their nutritional class (red=EAA, green=CEAA, blue=NEAA). (**d**) A model combining Dufton score and number of codons allocated to each amino acid was tested for its ability to predict liver protein expression levels.

**Fig 2c** illustrates the relative magnitudes of these properties for each amino acid (tabulated in full in **Supplementary File 1**). A MLR analysis of all properties applied to liver expression data indicated significant contributions from the number of codons allocated (p=5.2 × 10^−14^) and the Dufton score (p=5.2 × 10^−13^), but no significant influence of energetic cost. Combined in a single model, these two simple and immutable amino acid properties were sufficient to generate a moving average describing almost three orders of magnitude of liver expression (**Fig 2d**).

### Protective biological systems at extremes of amino acid composition

A prediction from these findings would be that animal proteins with the most extreme EAA or CEAA amino acid compositions would be the first to experience translational inefficiency in a state of amino acid deficiency. We theorised that such proteins might have retained counterintuitively extreme compositions to sense and respond to environmental adversity. When the fEAA and fCEAA composition of 20,397 human proteins was visualised (**Fig 3a**), outlier proteins were found to intersect with our understanding of the biology and pathology of animal survival during malnourishment.

**Figure 3.**
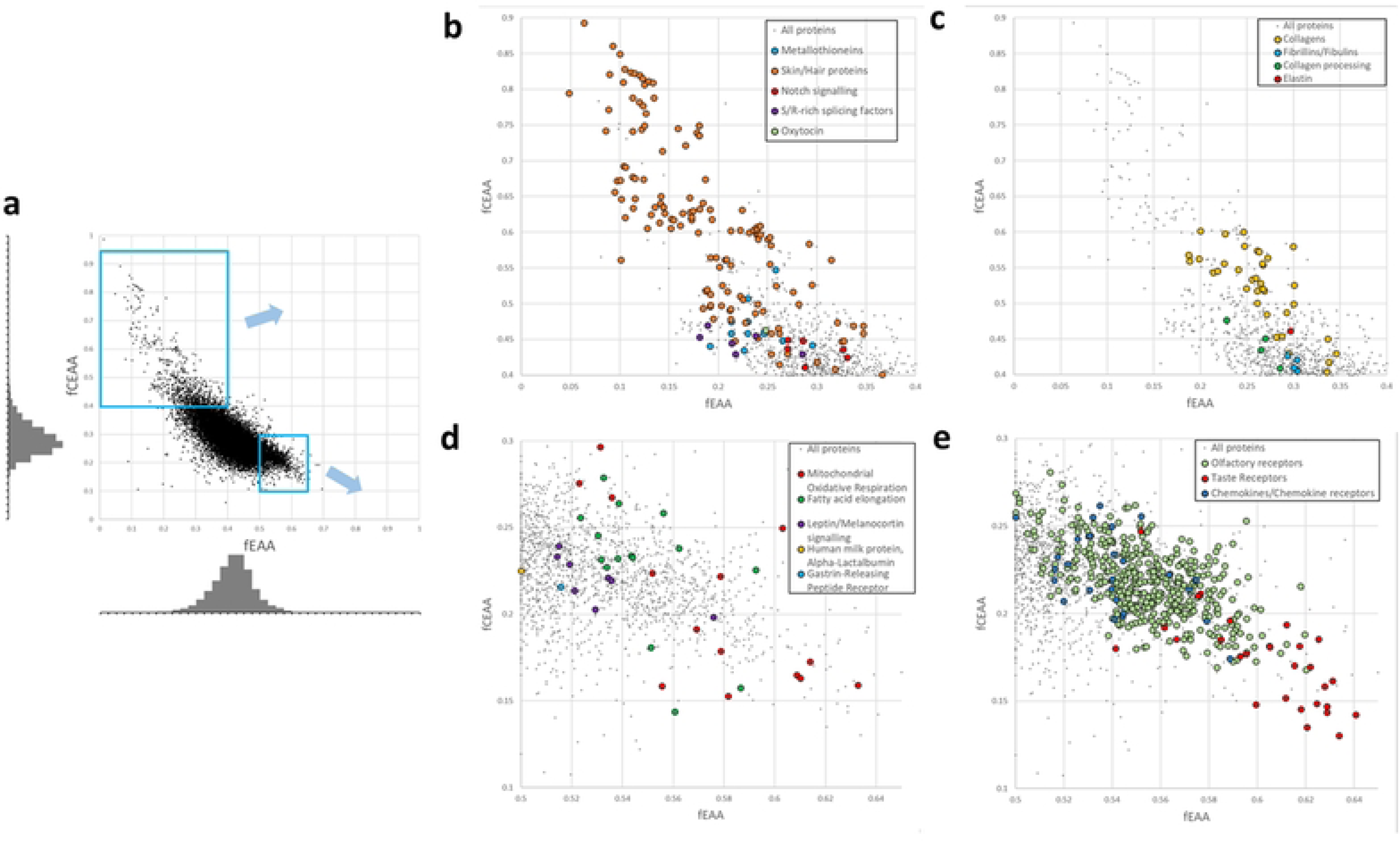
EAA and CEAA over-representation in specific proteins and pathways reveals the consequences of, and response to, amino acid deprivation. The composition of 20,397 human proteins was plotted, fEEA against fCEAA. (a) Tight distribution within a dense central cluster was clarified by the associated histograms. Magnified outlier sectors of a contained protein families or functionalities with extremely high fCEAA (b/c) and fEAA (d/e) likely susceptible to reduced expression during nutritional insufficiency.

The fCEAA outlier group primarily consisted of proteins with roles in the formation of connective tissue, skin, hair, and their maturation enzymes: collagens, elastin, keratin-associated proteins, late cornified envelope proteins, small cysteine, glycine and proline repeat-containing proteins, fibrillins, fibulins, lysyl oxidase, and latent-transforming growth factor beta-binding proteins (**Fig 3b** and **3c**)[24, 25]. Gastrointestinal excretion and the production of skin and hair are the three principal routes of irretrievable amino acid loss from the body [26], so the relative paucity of EAAs in these proteins may be a resource conservation strategy. Furthermore, their extreme CEAA composition may act as a translational regulator for life processes that can be temporarily sacrificed or permanently downscaled to conserve energy and amino acid reserves. Anorexia nervosa can be accompanied by thinning hair, brittle nails, and deterioration in skin health [27] and, separately, periods of bodily stress or illness are frequently recorded as discontinuous nail growth in the form of Beau’s lines or ‘pitting’. Other proteins with high fCEAA that might be susceptible to the effects of dietary deficiency include all 14 metallothioneins involved in heavy metal-binding and oxidative stress responses, several members of the serine-/arginine-rich splicing factor family, oxytocin (a hormone involved in all aspects of reproduction from sexual arousal, to uterine contraction in labour, mother-offspring bonding, and milk production) and multiple components of the Notch cell fate and differentiation signalling pathway (including DLL3, NOTCH4, and JAG2).

Extreme fEAA was observed in both leptin (LEP) and the melanocortin receptor proteins (MC1R-MC5R) (**Fig 3d**) – components of an established hypothalamic signalling pathway that responds to increased levels of adiposity by promoting satiety. This pathway is also linked to onset of puberty and stature [28]. GRPR (gastrin-releasing peptide receptor) also possesses high fEAA and similar appetite-suppressing role. We speculate that their conventional signalling pathways are augmented by ‘hard-wired’ protein synthesis inhibition during EAA deprivation – expression reduction of any of these components would act to increase appetite drive, potentially leading to recuperative ingestion of amino acids. Most members of the large olfactory receptor subfamily [29] exhibit high EAA density. Similarly, bitter taste receptors of the TAS2R subfamily have an extremely high EAA composition (**Fig.3e**) - with member TAS2R20 ranked 9^th^ in the entire proteome. This contrasts with the proteome-average EAA composition of the umami and sweet taste receptors of the TAS1R subfamily. The TAS2R family fulfilled an important survival function in human prehistory by allowing detection and rejection of potentially toxic substances in foraged food. We hypothesise that the proteins responsible for these two sensory modalities have evolved fragility of expression during dietary EAA deficiency. The resulting reduction in bitter taste and smell acuity may lower food discrimination or aversion, offering access to a greater range of foodstuffs potentially containing EAAs (histidine, tryptophan and valine salts are all bitter-tasting [30]). There is tentative evidence that prolonged anorexia nervosa blunts taste sensitivity [31]. Three other protein families have significant representation at the extremes of fEAA, including a group of fatty acid metabolism proteins (**fig 3d**), 14 protein components of the mitochondrial electron transport chain complexes (**fig 3d**) - and all 26 protein members of the CC-/CX-chemokine and chemokine receptor families that control chemotaxis and other immune cell functions (**Fig 3e**). Examination of the cell line data described earlier revealed globally reduced expression in the high fCEAA/high proliferation groups including the majority of the individual fCEAA outlier proteins discussed here. Highly proliferating lines showed specific reductions in expression of a number of fEAA outlier proteins, including most taste and olfactory receptors, some of the chemokines and their receptors, and slight decreases in expression for mitochondrial proteins such as MT-ND4, MT-ND5, and MT-ATP8.

*E. coli*, *S. cerevisiae*, and *A. thaliana* do not have compromised amino acid biosynthesis pathways requiring external EAA provision. However, they do possess the intrinsic ‘constraints’ defined in our earlier MLR findings as negative coefficient amino acids. We hypothesised that proteins at MLR-defined compositional extremes might also have been subject to evolutionary selection. Scores based on combined negative coefficients (MLR-) and combined positive coefficients (MLR+) were assigned to each protein based on composition. Proteins at the four MLR score distribution extremes were analysed via The Gene Ontology Resource [32] using Panther [33] to identify significant enrichment for specific biological processes (gene ontologies: GO). Next, MLR-scores and MLR+ scores were collated from the full set of proteins associated with identified gene ontology terms and statistically compared to the whole protein population using a two-tailed z-test. For *E. coli*, we observed significant enrichment within the less negative MLR-, more positive MLR+ protein sector for 251 proteins designated under the ‘*response to abiotic stimulus*’ GO term (MLR-p=2.6 × 10^−5^, MLR+ p=1.1 × 10^−6^). The two least negative MLR-proteins in the entire *E. coli* proteome fall within this environment-detection category: acid shock protein (asr) and cold shock-like protein (cspC). Also significant were 116 proteins under the ‘*translation*’ GO term (MLR-p=1.1 × 10^−18^, MLR+ p=9.8 × 10^−61^). This assessment is indirectly related to protein sequence and thus subject to bias from the presence of multiple paralogs. Reanalysis of the paralog-rich *translation* term, collapsing multiple paralogs to a single averaged archetype, still yielded significance (MLR-p=0.002, MLR+ p=5.5 × 10^−8^). *Translation*-related GO terms also presented statistically significant MLR score biases in *S. cerevisiae*, *A. thaliana* (root), and *H. sapiens* (liver). Likewise, environmental detection-related GO terms *response to high light intensity*, *water deprivation*, and *cold acclimation* (*A. thaliana*); *detoxification*, and *response to stress* (*H. sapiens*), all showed significant MLR score biases (**Supplementary File 2**).

### Amino acid composition and disease

In humans, gene-environment (GxE) interactions modifying disease risk and phenotypic expressivity may be encountered by proteins with extreme fEAA/fCEAA composition. Malnourishment in early life is currently experienced by 1 in 5 of the world’s population, affecting stature, intellectual ability, future fertility, and risk of chronic illnesses – with the WHO reporting 145 million children with stunted height in 2020 [34, 35]. In the first approach to examine potential EAA/CEAA deficiency influences on disease risk, the online DisGeNET catalogue of genes associated with 8,383 diseases [36] was queried to identify extreme fEAA/fCEAA composition proteins which also had robust aetiopathological roles supported by at least 10 distinct disease annotations (**Supplementary File 1**). Proteins linked to cancers (e.g., the tumour suppressor, CDKN2A), CNS disorders (e.g., the myelin constituent, PMP22), and developmental disorders (e.g., skeleton and tooth development protein, SLC10A7) were represented at fCEAA and fEAA extremes. For fCEAA, there were many connective tissue disorders (due to the collagen protein family), as well as several proteins linked to miscarriage (COL5A1, IGFBP6, LGALS3); for fEAA, proteins were linked to immunological disorders, primarily due to the chemokine family and their receptors.

In a second, multigenic approach, five conditions (cancer, male infertility, female infertility, tooth abnormality, obesity) and one phenotype (stature/height) were chosen as established indicators of malnourishment or, in the case of cancer, selected because of a pathology defined by aberrant proliferation. Risk proteins for each disorder were compiled from the literature or public databases and two-tailed Z-tests performed to determine if risk protein lists showed average fCEAA or fEAA values significantly deviating from the entire proteome (**Supplementary File 1**). Significant findings were observed for cancer and stature. Established cancer proteins (n=723, from the COSMIC Cancer Gene Census [37]) showed a highly significant under-representation of EAA (p=0.00 × 10^0^) and compensatory increase in fNEAA (p=2.23 × 10^−31^). Stature genes (n=116, manually curated from literature) showed an increase in fCEAA (p=1.02 × 10^−6^) and a decrease in fEAA (p=1.02 × 10^−05^) although significance was largely driven by 8 members of the collagen family.

## Discussion

We have established both extrinsic and intrinsic rules by which a protein’s amino acid composition can influence its expression. The profile of extrinsic CEAA inhibitory effects on expression during baseline and proliferative cellular conditions offers the first rigorous molecular definition of this historically ignored nutritional class. One consequence of our findings is that the proliferative state of laboratory cell lines (largely unreported in publication methods) may be a confounding factor for experimental replication of functional or expression studies - for half of all proteins. Expression of proliferation biomarker MKI67 may be a useful benchmark for such studies. By contrast, only modest consequences were observed for EAA-enriched proteins. Determining the true scale of extrinsic EAA influences on protein expression *in vitro* and *in vivo* will require experimentation with amino acid-deficient culture media/feeds. The remarkable second finding that individual amino acids affect protein expression in a largely conserved manner across species appears to be a consequence of the intrinsic amino acid properties of size, structural complexity, and codon allocation. It is presumed that these intrinsic effects act at the ribosome during translation. We observed that proteins participating in translation and in species-specific environmental stress responses were significantly under-represented in amino acids with negative influence on expression and, conversely, over-represented in those with positive influence. We suggest this selective drive has ensured that survival-enhancing proteins can be rapidly, robustly, and highly expressed, even in challenging cellular and environmental conditions. Human and animal proteins at the extremes of EAA/CEAA composition may also have evolved as an advantageous strategy to survive extrinsic nutritional scarcity. As well as the described effects on hair/nail/skin production and food-seeking behaviours, the modulation of collagen protein expression may be a key response to nutritional status in development: determining the limits of body size and appropriate maternal resource allocation - and aligning future metabolic demand (proportional to body scale) with anticipated environmental resource availability.

The findings presented here offer public health programs the prospect of quantitative protein biomarkers of clinical and sub-clinical amino acid dietary insufficiency. Additionally, they inform current clinical interventions based on amino acid supplementation and restriction. There are established benefits to amino acid supplementation (primarily the CEAAs glutamine or arginine) for patients undergoing ulcer treatment or post-surgery wound healing [38–41], potentially promoting connective tissue proliferation/regeneration. Supplementation with other CEAAs such as cysteine, or negative MLR coefficient amino acids such as serine, isoleucine, and tryptophan are now also worthy of investigation. By contrast, restricting amino acids in diet is an emerging concept in cancer treatment, capitalising on the specific demands of tumour cells. Our earlier findings on proliferation demands suggested that this ‘hallmark’ [42] would manifest as reduced fCEAA in cancer risk proteins. In fact, cancer proteins exhibited an extraordinary under-representation of EAA, suggesting that restricted essential amino acid supply to the tumour microenvironment may be a major determinant of protein expression, genotype-phenotype correlation, and clonal selection in cancer [43]. In tumours, expression of high fEAA proteins involved in mitochondrial oxidative respiration and chemokine function may thus be compromised. This would be consistent with the Warburg effect [42] which describes the metabolic shift within tumours from oxidative respiration to glycolysis, and it may also contribute to the extensive chemokine/receptor-mediated interactions between tumour cells, stromal cells and macrophages [44]. Multiple amino acids have been trialled in restriction studies [45], mostly on the basis of gross abundance, so the detailed findings reported here may guide future dietary protocols in cancer treatment.

## Methods

### Data import and basic amino acid frequency analysis

From Uniprot.org, one representative human protein sequence per gene (totalling 20,397) was downloaded from the Reference Human proteome (ID: UP000005640) in FASTA format. Microsoft Excel text analysis formulas were applied to calculate the total amino residues, the frequency of each individual amino acid, and the relative proportions of the EAA/CEAA/NEAA nutritional classes present within each protein (**Supplementary File 1**). For example, a total of 34 EAA amino acids within a protein of 299 residues generates a fEAA of 0.11. Frequencies of amino acids or amino acid classes were used to remove the confounder of protein length differences. Similar processes were carried out for the *Escherichia coli* (UP000635675), *Saccharomyces cerevisiae* (UP000002311), and *Arabidopsis thaliana* (UP000006548) proteomes.

### Protein expression correlation with amino acid classification

Protein expression data were imported as simple .txt files into Excel from publicly available datasets in PaxDB. Species-specific protein identifiers in the expression data were converted into universal UniProt or UniPARC identifiers using the VLOOKUP command accessing imported conversion tables, allowing correlation with the amino acid/amino acid class frequencies of each protein.

### Moving median/average expression analysis

Nutritional amino acid class frequency was ranked and plotted against the moving median liver protein expression level (**Fig.1a**, periodicity of 469 = 5% of total). Multiple linear regression (see below) model score for each protein was plotted against liver expression value (log scale) and an Excel moving average trendline applied (periodicity of 100 proteins) (**Fig. 2a/b/d**).

### Tissue and cell line plots of changing EAA/CEAA/NEAA proportions across expression levels

For a tissue or cell line, both the numerical expression levels and the fEAA, fCEAA, and fNEAA for each protein were separately converted into ranks. Cumulative average proportions for each amino acid class were calculated from lowest-to-highest expressed proteins and, in parallel, highest-to-lowest expressed proteins – the average of the pair of values was calculated for each individual protein and plotted (**Fig.1b-g, Supplementary Figure 1, Supplementary File 1**). This method produced smoothed plots of trends in relative amino acid class representation as a function of ranked protein expression level.

### Multiple Linear Regression and models

Multiple linear regression (MLR) in the Excel ‘Data Analysis’ ToolPak add-in was used to identify intrinsic parameters or individual amino acids with significant correlation to protein expression level, and their respective coefficients (**Supplementary File 1**). Statistically significant (p<0.05) MLR coefficients were normalised across organs and species in **Table 1**. MLR findings allowed the construction of models which generated relative numerical expression predictions for each protein based on coefficients and amino acid frequencies.

### Statistical tests of gene ontology and disease candidate lists

Two-tailed z-tests were applied to groupings derived from MLR+ (combined coefficients increasing expression) and MLR- (combined coefficients decreasing expression) data, or fEAA/fCEAA/fNEAA data. For human fEAA and fCEAA, analysis of parent population distributions showed statistically significant deviations in kurtosis and skewness. However, the large population size and small Lilliefors D effect size values of 0.043 (fEAA) and 0.078 (fCEAA) justified treating the distributions as effectively normal for the purpose of z-tests (**Supplementary File 1**). Full monogenic and multigenic disease lists and statistical analyses relating to nutritional class composition are found in **Supplementary File 1**. Statistical analysis of enriched GO terms in MLR- and MLR+ data are found in **Supplementary File 2**.

## Acknowledgements

R.T. was funded through The Robertson Trust Internship Scheme 2021

## Data sharing

The human proteome data and majority of other analyses are to be found in the Supplementary Information files.

---

***Supplementary Figure 1 The relative proportions of the three nutritional amino acid classes change with protein expression level and proliferation rate (extended data from Fig.1b-g).** A smoothing procedure (see **Methods** and **Supplementary File 1**) was applied to visualise trends in relative, ranked amino acid class proportion when plotted against ranked protein expression level for 8 human tissues and two samples of a lung cancer cell line, A549 (data from PaxDB). As described in the main text, tissues can be placed in three groups (**a, b**, and **c**) based on the profile of EAA/CEAA/NEAA composition across the range of expression levels. A549 differences (**d**) most likely represent amino acid constraint effects brought about by different proliferation rates. Graph **e** shows the same liver data as in **a** but with randomised expression level as a control.*

***Supplementary File 1:*** *Multiple datasets and analyses comprising; Amino acid composition calculator, Human proteome AA composition, 25 selenocysteine-containing proteins, Testing populations for normal distribution, Calculating and smoothing relative proportions of EAA, CEAA, and NEAA in proteins as a function of expression level, Multiple Linear Regression (MLR) analyses across organs, species, and amino acid parameters, Extreme fEAA-& fCEAA-associated diseases, and Multigenic disease statistics*.

***Supplementary File 2:*** *Statistics of the over-represented Gene Ontologies (GOs) in proteins with extremes of MLR coefficients*.

